# A New Mathematical Model for LVAD-Supported Ventricles: Direct Parameterization from Ramp-Test Clinical Data and Verification via Hybrid Modeling

**DOI:** 10.64898/2026.03.20.712251

**Authors:** E. Umo Abraham, A. Welch Brett, Arman Kilic, O. Kung Ethan

## Abstract

**Background:** Conventional left ventricular assist device ramp metrics are load dependent, obscuring intrinsic myocardial recovery. A mechanistic, patient-specific representation of ventricular mechanics, identifiable from routine clinical data, could provide quantitative indices of intrinsic left ventricular (LV) function for longitudinal recovery surveillance.

**Objective:** To develop and verify a ramp-integrated, patient-specific model of HeartMate 3-assisted LV function that can yield indices of intrinsic myocardial contractility and remodeling.

**Methods:** We represented LV pressure–volume (PV) behavior with a PV envelope composed of a monotonic passive PV relation (pPVR) and a unimodal active PV relation (aPVR). We developed a parameterization procedure to infer the patient-specific shape of this envelope directly from routine ramp-test data. We then embedded the parameterized envelope within the PSCOPE framework, a hybrid platform that couples a lumped-parameter network to a physical HeartMate 3 flow loop, to reproduce clinical ramp hemodynamics. Percent residuals between simulated outputs and the corresponding clinical measurements verified the implementation of the PV envelope within PSCOPE.

**Results:** In three HeartMate 3 recipients, the PSCOPE models reproduced ramp hemodynamics with residuals generally ≤ 20% across pump speeds and measured variables. Cardiac index residuals ranged from 0–18.5%, systemic and pulmonary arterial pressure residuals remained ≤ 18.4%, and pulmonary arterial wedge pressure residuals remained ≤ 20%. The PSCOPE models matched central venous pressure within ≤ 3 mmHg in all cases, although one setting yielded a 33.3% residual due to a low reference pressure. For one patient, the model reproduced ramp hemodynamics at a speed deliberately withheld from PV-envelope parameterization with residuals ≤ 10%, supporting cross-speed generalizability. Patient-specific PV envelopes also revealed clinically meaningful heterogeneity in LV diastolic stiffness, volume threshold for declining systolic function, operating PV points for peak systolic function, and contractile reserve.

**Conclusions:** Ramp-integrated parameterization of the monotonic pPVR and unimodal aPVR yields a compact, mechanistic PV envelope that is identifiable from routine clinical data and verifiable within PSCOPE. The resulting indices characterize intrinsic LV function and may enhance longitudinal recovery surveillance and inform LVAD management. Prospective multicenter validation is warranted to confirm the generalizability and clinical utility of this approach.

## 1. Introduction

Heart failure remains a pervasive and challenging condition affecting millions worldwide [1]. Although heart transplantation remains the gold standard therapy for end-stage heart failure, donor scarcity and patient contraindications limit access [2]. Consequently, mechanical circulatory support has become a critical option for advanced cases. Among mechanical circulatory support modalities, left ventricular assist devices (LVADs) provide durable systemic support for patients with left ventricular (LV) cardiomyopathy. Clinical strategies for LVAD management include destination therapy, bridge-to-transplant, and bridge-to-recovery. The bridge-to-recovery strategy is particularly compelling because it aims at substantive myocardial recovery [3-4]. This pathway is viable only when LVAD-mediated unloading induces LV reverse remodeling—reduced chamber size, more favorable geometry, and improved systolic function—such that in selected patients, recovery is sufficient for LVAD explantation or decommissioning, as shown in multicenter programs [5].

LV myocardial contractility, the ability to generate pressure and eject blood, is fundamental to the pathophysiology of heart failure [6]. Progressive declines in contractile function, arising from myocyte injury, ischemia, or sustained volume/pressure overload, drive adverse remodeling and disease progression [7]. Conversely, LVAD support can promote reverse remodeling with concomitant improvements in LV contractility, signifying a potential trajectory towards recovery [8-10]. Clinicians typically assess remodeling and contractile reserve using echocardiography and hemodynamic measurements acquired during incremental LVAD speed adjustments (“ramp” testing), which inform assessments of recovery potential and explant candidacy [11–13]. Many programs interpret these assessments using threshold criteria for cardiac index (CI), left-ventricular end-diastolic diameter (LVEDD), ejection fraction, mitral regurgitation, aortic valve opening, pulmonary arterial pressure (PAP), and systemic arterial pressure (AP) [14–16]. A central limitation of this approach is load dependence; these metrics are highly sensitive to changes in preload and afterload. Stepwise speed changes during ramp testing transiently alter loading conditions and may confound interpretation, making the same ventricle appear “improved” or “deteriorated” without a corresponding change in intrinsic contractile function [17–18]. This interpretive confounding, compounded by heterogeneity in ramp protocols, likely contributes to the wide range of reported post-explant recovery rates (2%–73%) across programs [19–21].

This work introduces the Ramp-Derived Elastance formulation (RDEF), a novel empirical model of the pressure-volume (PV) dynamics in the failing ventricle that can be personalized to a patient using routine ramp-test data. Rather than interpreting LV function directly from load-dependent ramp outputs, RDEF uses ramp-derived operating points to estimate an underlying passive PV relation (pPVR) and active PV relation (aPVR) that govern the ventricle’s preload and afterload sensitivities. The resulting RDEF parameters enable the derivation of indices that are sensitive to myocardial remodeling and agnostic to acute changes in preload and afterload induced by stepwise LVAD speed changes. We verify the RDEF model within the PSCOPE hybrid cardiovascular framework [22–25] by developing digital twins for three HeartMate 3 (HM3) recipients and comparing simulated hemodynamics with contemporaneous ramp measurements. RDEF yields mechanistic, noninvasive indices of LVAD-assisted intrinsic LV function that complement standard echocardiographic and hemodynamic assessments. Clinical integration of these indices could reduce reliance on load-dependent ramp metrics, standardize longitudinal surveillance of LV recovery, and refine patient selection for LVAD explantation or decommissioning.

## 2. Methods

### 2.1 Study Design and Cohort

We conducted a retrospective single-center study to derive and clinically verify the RDEF model of LV failure. Twenty adults with HM3 support who underwent clinical ramp testing at the Medical University of South Carolina were screened. Inclusion criteria were: (i) ≥ 3 discrete LVAD speed settings within a single session; (ii) availability of ramp hemodynamics including heart rate (HR), CI, AP, PAP, pulmonary arterial wedge pressure (PAWP), and central venous pressure (CVP); and (iii) echocardiography adequate to compute LV end-diastolic volume (LVEDV) at each speed. Three patients met these criteria (Table 1).

**Table 1.**
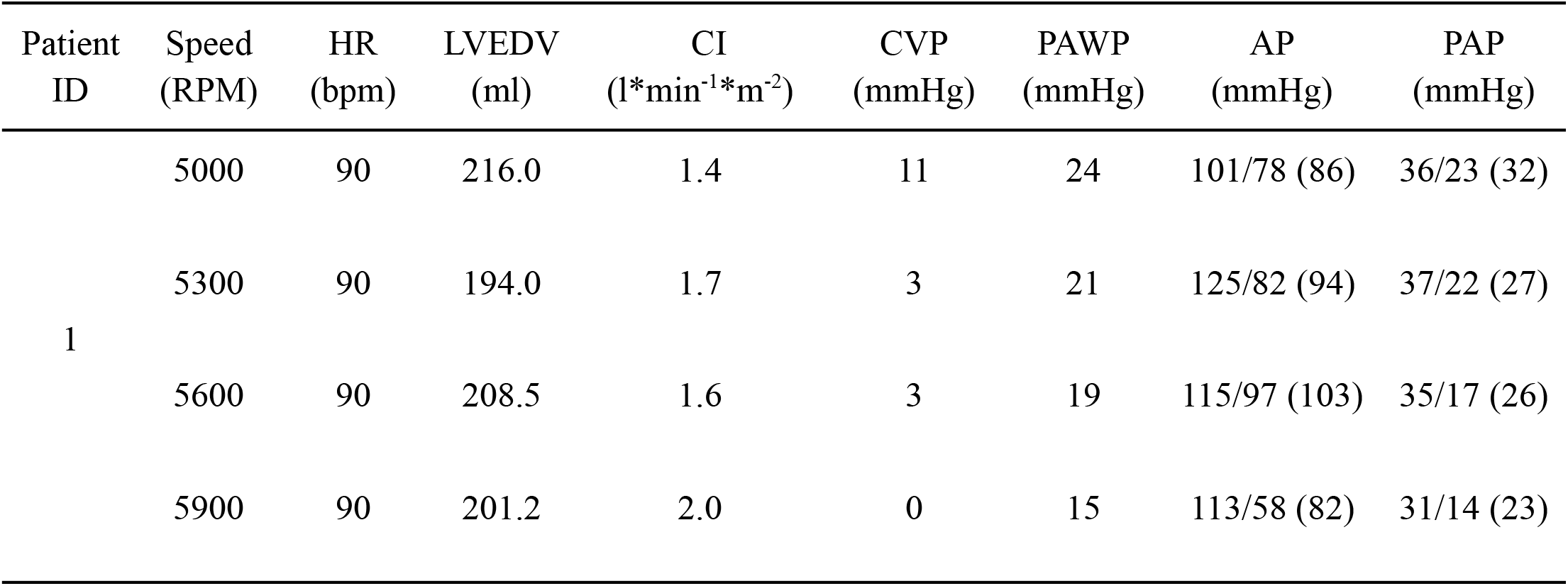
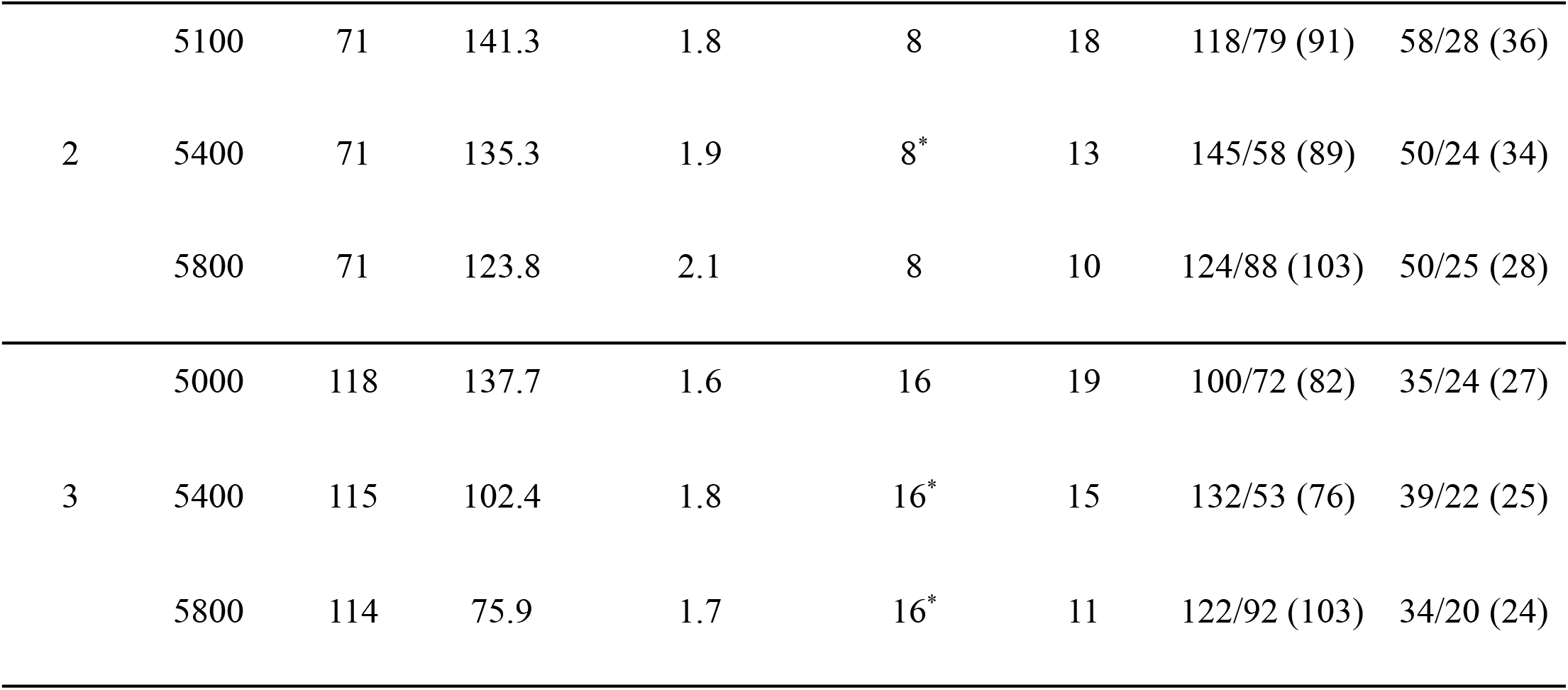
Summary of clinical ramp hemodynamics for the HeartMate 3 patient cohort. HR—heart rate; CI—cardiac index; AP—systemic arterial pressure; PAP—pulmonary arterial pressure; PAWP—pulmonary arterial wedge pressure; CVP—central venous pressure; LVEDV—left-ventricular end-diastolic volume; RPM—revolutions per minute. AP and PAP values are reported in “systolic/diastolic (mean)” format. “*” indicates values that were imputed due to missing clinical data.

CVP was unavailable for patient 2 at 5,400 RPM and for patient 3 at 5,400 and 5,800 RPM. In these instances, CVP was imputed by carrying forward the nearest ramp-acquired value within the same session, consistent with prior reports of CVP stability during HM3 ramp testing [26-27].

The Institutional Review Board of the Medical University of South Carolina approved this study. Given the retrospective design and use of de-identified data, informed consent was waived.

### 2.2 RDEF Formulation of LV Failure

RDEF models PV behavior in the failing LV using two relations: (i) a monotonic pPVR (Eq. 1), and (ii) a unimodal aPVR (Eq. 2) that reproduces the ascending and descending Frank–Starling limbs observed in failing ventricles [28–30] (Fig. 1). These relations are combined via a weighted superposition to compute instantaneous PV dynamics across the cardiac cycle (Eq. 3).

**Figure 1.**
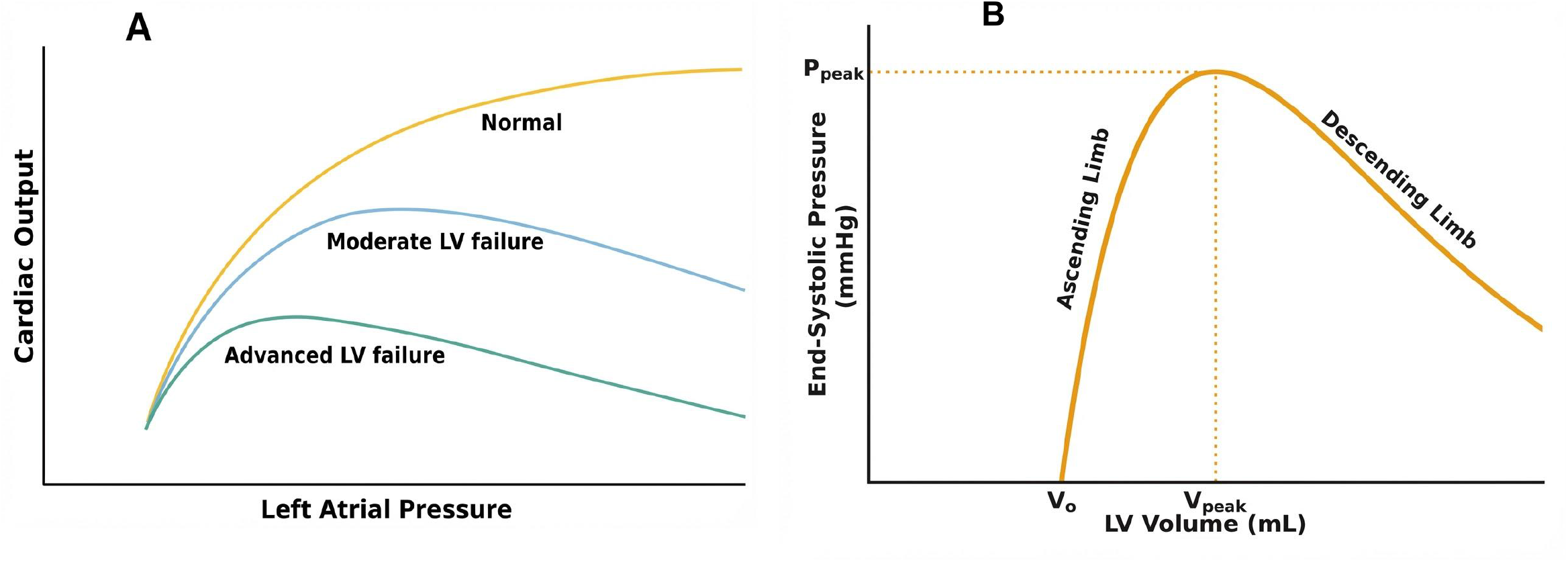
(A) Representation of the Frank-Starling curve for normal and LV failure conditions (adapted from [30]). (B) Representation of the unimodal aPVR capturing ascending and descending Frank-Starling limbs.

#### 2.2.1 Passive Mechanics: Monotonic pPVR

RDEF represents passive LV mechanics using the standard exponential pPVR [31-33]:

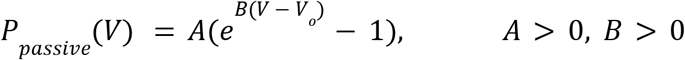

Where: *P*_*passive*_= Passive LV Pressure

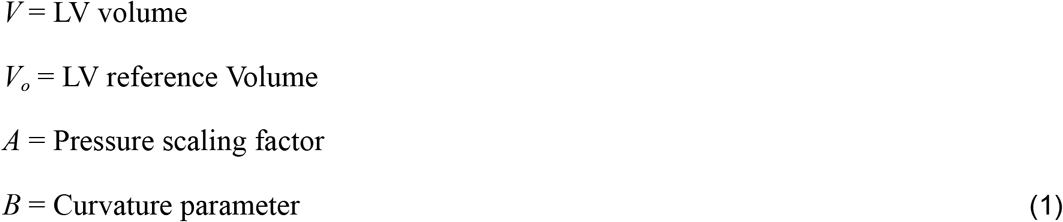

This expression is strictly monotonic with *P*_*active*_ (*V*_*o*_ ) = 0 and 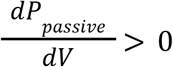 for all *V*≥*V*_*o*_.

#### 2.2.2 Active Mechanics: Unimodal aPVR

RDEF represents active LV mechanics with a unimodal aPVR:

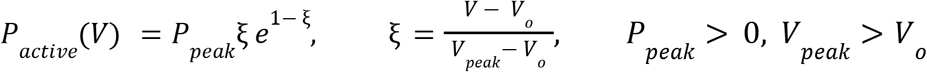

Where: *P*_*active*_= Active LV Pressure; i.e., pressure at maximum contraction

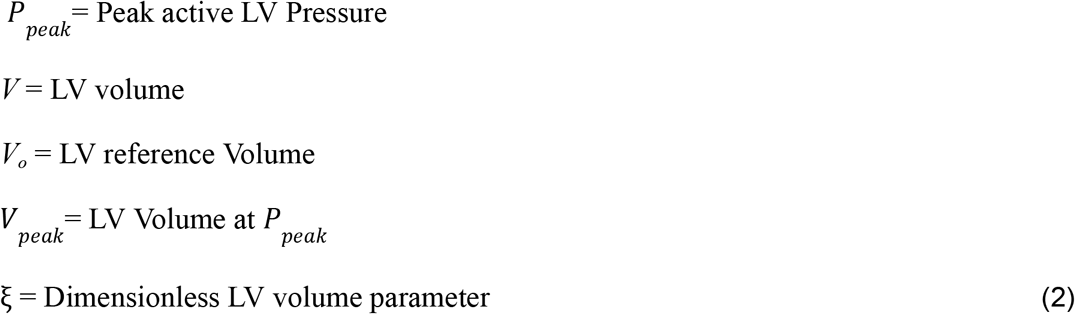

This expression is strictly unimodal: *P*_*active*_ increases for ξ < 1, attains a maximum at ξ = 0, and decreases for ξ > 1. The coordinate (*V*_*peak*_, *P*_*peak*_) specifies the operating point of peak systolic function (maximal pressure generation) and demarcates the region of increasing systolic function (ascending limb) from that of declining systolic function (descending limb) [34].

#### 2.2.3 Instantaneous PV Mechanics

A standard active–passive ventricular formulation computes instantaneous LV pressure over the cardiac cycle via a superposition of the passive and active PV components. Following Vasudevan et al. [28], a normalized double-hill function, ϕ(*t*), activates the aPVR (Eq. 1) and superimposes the pPVR (Eq. 2) via a weighted summation:

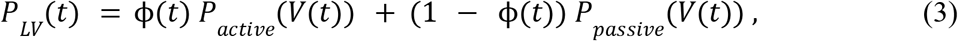

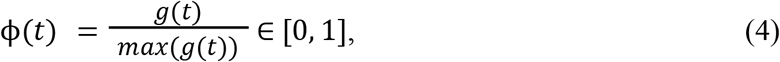

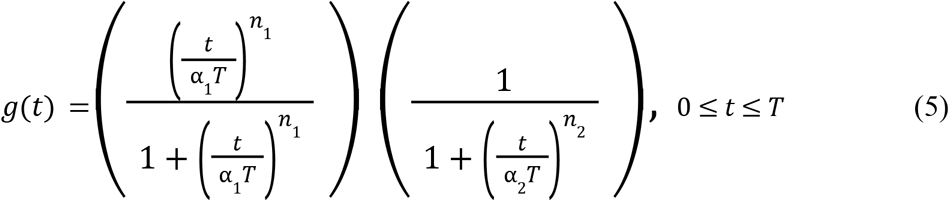

Where: *P*_*LV*_ (*t*) = Instantaneous LV pressure

t = Time point in the cardiac cycle

T = Cardiac period

A_1_, α_2_ = Time scales governing systolic and diastolic phases

*n*_1_, *n*_2_ = Dimensionless shape exponents

#### 2.2.4 RDEF Parameterization

Clinically indicated HM3 ramp testing yielded speed-specific hemodynamic and echocardiographic profiles for each patient (Table 1). At each pump speed, we computed LVEDV from echocardiographic LVEDD using the Teichholz relation [35]:

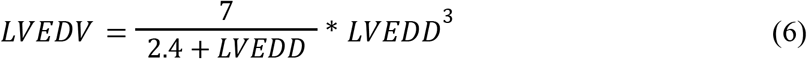

Acquiring true PV loops of the LV under active LVAD support is rarely feasible because it requires invasive LV catheterization and substantial reduction of LVAD support to reveal native LV pressure generation, which presents significant hemodynamic risk in this population. Therefore, we leverage ramp hemodynamics to approximate in vivo systolic and diastolic PV dynamics at each pump speed. For each speed in the ramp protocol, we combine echocardiography-derived LVEDV with contemporaneous systemic and pulmonary pressure measurements to define practical surrogates for invasive PV measurements of the LV. The ramp echocardiography dataset did not include LV end-systolic diameter (LVESD) and therefore did not support the reliable derivation of LV end-systolic volume (LVESV). Under continuous flow HM3 support, particularly at higher speeds with reduced pulsatility and frequent aortic valve closure, the LV often exhibits minimal native ejection with limited volume changes between end-diastole and end-systole. Under these conditions, LVEDV provides a reasonable approximation of the chamber volume throughout the cardiac cycle. Consequently, we used LVEDV as a pragmatic volume measurement when defining both the diastolic and systolic operating points. Specifically, we paired LVEDV with systolic-AP to form a systolic operating point (LVEDV, systolic-AP) and with PAWP to form a diastolic operating point (LVEDV, PAWP). Collectively, these speed-specific surrogates provide the constraints for patient-specific fitting of the empirical PV relations without LV catheterization.

The set of (LVEDV, systolic-AP) coordinates obtained within a ramp session was implemented as targets to estimate {*P*_*peak*_, *V*_*peak*_, *and V*_*o*_} by bound-constrained least squares optimization:

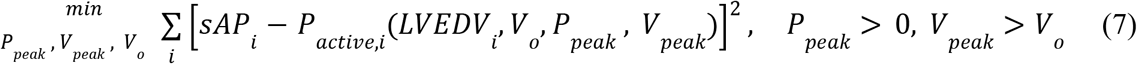

Where subscript “i” denotes a speed-specific parameter, and sAP denotes systolic arterial pressure. The fitted aPVR intersects the volume axis at a unique zero-pressure volume (*V*_*o*_, 0). We use this *V*_*o*_ as the reference volume in the pPVR so that both active and passive PV relations share a single zero-pressure reference point.

With *V*_*o*_ fixed from the fitted aPVR, the speed-specific (LVEDV, PAWP) pairs from the same ramp session were used to estimate {*A, B*} for the exponential pPVR via a second bound-constrained least-squares optimization:

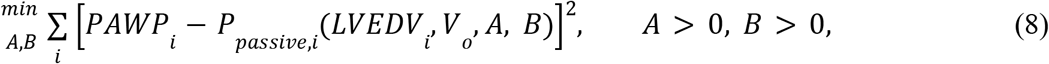

This protocol estimates patient-specific parameter values {*P*_*peak*_, *V*_*peak*_, *V*_*o*_, *A, B*} that personalize the RDEF model, yielding a compact, mechanistic summary of intrinsic LV behavior (Table 2). The personalized RDEF uses ramp-derived surrogate operating points to construct a continuous representation of LV PV dynamics. For each patient, three pump-speed settings were used for parameter identification, and the least-squares fitting relied exclusively on LVEDV, PAWP, and sAP at those speeds. Consequently, other contemporaneous hemodynamic measurements (CI, mean/diastolic AP and PAP, CVP) played no direct role in fitting the PV relations and instead served as independent quantities for assessing whether PSCOPE digital twins embedded with the personalized RDEF could reproduce the full ramp hemodynamic profile. For patient 1, the 5,300 RPM condition was additionally withheld entirely from the parameter identification process and used solely to evaluate the digital twin’s performance at a pump speed and loading condition not used for personalizing the RDEF model.

**Table 2.** Patient-specific LV module parameters. The unimodal aPVR is parameterized by {*P*_*peak*_, *V*_*peak*_, *V*_*o*_} and the monotonic pPVR by {*A, B*}. Parameters are held fixed across all speed-specific digital twins for the same patient.

| Patient ID | $P_{peak}$ (mmHg) | $V_{peak}$ (ml) | $V_o$ (ml) | $A$ (mmHg) | $B$ (ml<sup>-1</sup>) |
| --- | --- | --- | --- | --- | --- |
| 1 | 117.7 | 205.2 | 188.9 | 23678.1 | 4.0E-05 |
| 2 | 156.3 | 130.1 | 118.3 | 21550.0 | 3.8E-05 |
| 3 | 134.4 | 93.1 | 47.6 | 48200.0 | 5.0E-06 |

### 2.3 PSCOPE Digital Twin Verification

We verified the RDEF model by integrating it into PSCOPE, a hybrid cardiovascular platform that couples a lumped-parameter network (LPN) to a physical HM3 benchtop circuit via an iterative flow–pressure coupling algorithm [22–25]. In this study, a “digital twin” refers to a patient-specific PSCOPE configuration that reproduces clinical ramp hemodynamics across varying LVAD speed settings. We constructed these digital twins by embedding each patient’s personalized RDEF as the LV component within the LPN and synchronizing the in-vitro HM3 speed setting with the clinical ramp protocol.

Our verification aims to test the hypothesis that the RDEF remains invariant across pump speeds, as it characterizes the patient’s intrinsic myocardial state. Conversely, we treat each ramp speed as a distinct physiologic state to capture acute cardiovascular adaptations induced by ramp-unloading of the ventricle. By maintaining constant RDEF parameters while updating relevant LPN inputs to simulate ramp-induced cardiovascular adaptations, we test the digital twin’s ability to reproduce speed-specific hemodynamics. Successful reproduction of the entire clinical ramp protocol under a fixed RDEF affirms that our formulation effectively decouples intrinsic cardiac function from the confounding influences of pump speed and loading conditions. We quantified the digital twin’s performance using the percent residuals between simulated hemodynamic outputs and the corresponding clinical measurements.

The LPN used in this study was adopted from our previous work (Fig. 2A) [22]. At each ramp speed, we prescribe HR, systemic vascular resistance (SVR), and pulmonary vascular resistance (PVR) in the LPN according to clinical ramp measurements. The LPN accepts HR as a direct input to set the simulated cardiac period. Systemic and pulmonary resistance parameters were initialized according to previously reported reference values [36] and then scaled uniformly using distinct scaling factors for their respective resistance networks to match clinical SVR and PVR targets. This approach preserves the reference distribution of proximal versus distal resistances in simulated vessels while enforcing the prescribed aggregate SVR and PVR at each speed.

**Figure 2.**
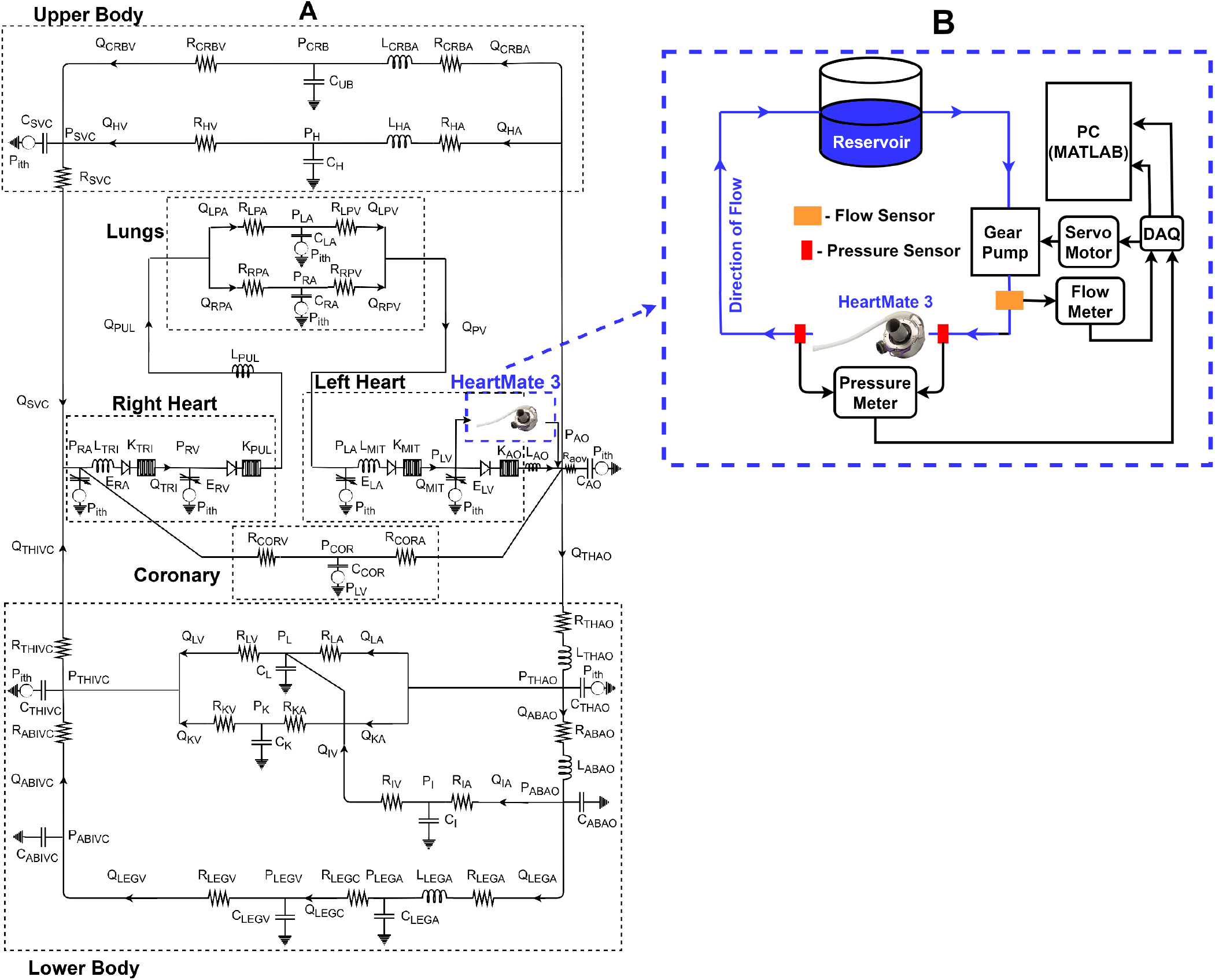
PSCOPE hybrid digital-twin implementation (adapted from [22]). (A) Patient-specific LPN embedded with personalized RDEF parameters to simulate ramp hemodynamics. (B) Benchtop HM3 flow loop replicating ramp unloading of LV.

With these primary clinical inputs fixed, we systematically tuned the remaining LPN parameters as per our established protocols [22] to optimize the digital twin’s replication of the patient’s baseline physiologic state (typically the speed closest to the patient’s resting clinical state). These baseline LPN input values are used to initialize simulations for all other ramp speeds. For subsequent speed settings, we tune only a restricted subset of LPN parameters, specifically circulating blood volume, aortic capacitance, and pulmonary capacitance, to minimize residuals between simulated and measured hemodynamics. We selected this specific subset to represent autoregulatory responses such as changes in aortic compliance, pulmonary arterial compliance, and fluid distribution, which are likely to occur during ramp-unloading of the LV. Conversely, LPN parameters governing peripheral vascular compliance, inertance, and cardiac chamber properties were held constant, as these are unlikely to undergo acute changes during the limited timeframe of a ramp test.

In specific cases, we incorporated targeted speed-dependent updates of specific LPN parameters to reflect documented pathological changes rather than generalized autoregulation. For Patient 1 at 5,000 RPM, a documented positive Kussmaul’s sign (a paradoxical rise in jugular venous pressure during inspiration indicating impaired right ventricular filling) [37-38], motivated the tuning of right ventricular diastolic elastance. For Patient 3, clinically documented worsening of tricuspid regurgitation at higher speeds necessitated speed-specific tuning of the tricuspid valve flow parameters. These targeted adjustments enabled the digital twins to capture the distinct pathological changes documented in specific patients.

Consistent with our prior validation study [22], PSCOPE couples the LPN to a benchtop HM3 flow loop to enable closed-loop modeling of LVAD-supported hemodynamics (Fig. 2). At each coupling iteration, the hybrid framework prescribes inflow conditions to the HM3, measures the resulting pump pressure head, and imposes that pressure head as a boundary condition across the LPN segments adjacent to the LVAD’s inlet and outlet. Iterations proceed until experimental and simulated HM3 flows converge (target NRMSE ≤ 20%) [22].

## 3. Results

### 3.1 PV Envelope

The fitted pPVR and aPVR intersect to define patient-specific PV envelopes (Table 2; Fig. 3), which support the mechanistic interpretation of intrinsic native LV properties. The geometry of the PV envelope characterizes the diastolic stiffness and contractile capacity of native LV function. The passive elastance, defined as the local slope of the pPVR [39], indexes diastolic stiffness (which is a function of the filling volume):

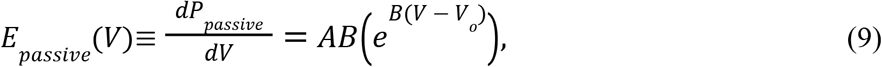

**Figure 3.**
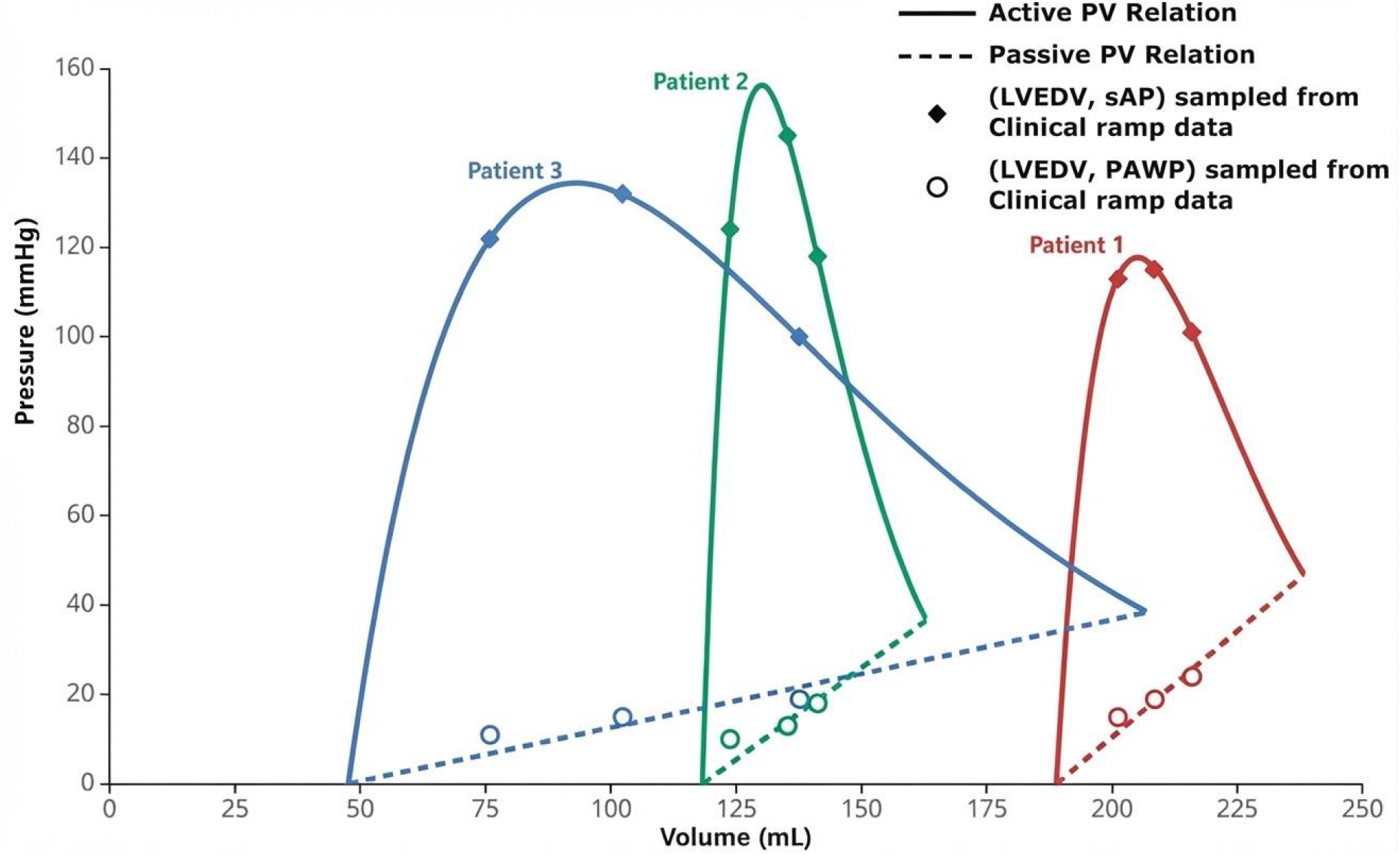
Patient-specific PV envelopes under HM3 support. Systolic (LVEDV, sAP) and diastolic (LVEDV, PAWP) surrogate operating points for the LV are also shown.

Larger *E*_*passive*_(*V*) corresponds to a steeper passive curve and a stiffer chamber. We evaluate *E*_*passive*_(*V*_*peak*_ ) to characterize maximal LV stiffness before the onset of chronic volume overload (*V* > V_*peak*_).

To quantify contractile capacity, we adopt an energy-based approach inspired by the PV area (PVA) framework developed by Suga [40]. PVA quantifies the total mechanical energy produced by the ventricle over a cardiac cycle and may be evaluated under different contractile conditions. For an idealized fully isovolumic beat, in which the ventricle undergoes an entire cardiac cycle at fixed chamber volume *V* (i.e., without ejection or filling), the PVA is defined as the area bounded by the active and passive PV relations over the volume interval [*V*_*o*_, *V*] [41–43]. Although isovolumic contraction produces no ejection and thus no external stroke work, the ventricle still expends mechanical energy as stored (potential) energy while generating pressure at constant volume.

We denote this isovolumic contractile energy as *PVA*_*iso*_, interpreted as the maximal contractile energy available at a specified preload *V*. We can derive *PVA*_*iso*_ from the PV envelope for a given preload *V* as:

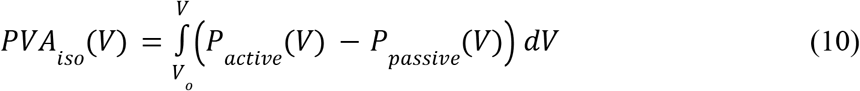

i.e., the area between the aPVR and pPVR from *V*_*o*_ to *V*. Evaluating this quantity at *V*_*peak*_ and normalizing by body-surface area (BSA) gives,

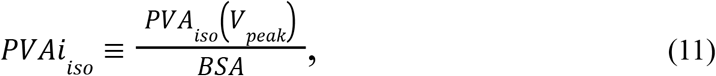

which serves as an energetic index of contractile reserve at peak systolic function [44], before the ventricle operates in the descending Frank-Starling limb associated with declining systolic function. Therefore, larger *PVAi*_*is*o_ values indicate a greater capacity to sustain effective systolic function across changing loading states.

#### 3.1.1 Patient Cohort

The PV envelopes derived for this cohort reveal marked inter-patient variation in LV size, diastolic stiffness, and contractile reserve (Fig. 3). Patient 1 exhibited the most dilated and stiff ventricle (*V*_*o*_ = 188.9 ml; *E*_*passive*_(*V*_*peak*_) = 0.95 mmHg*ml^-1^) with the lowest contractile reserve (*P*_*peak*_ = 117.7 mmHg; *PVAi*_*is*o_ = 585.2 mmHg*ml*m^-2^). Patient 2 had the highest peak active pressure (*P*_*peak*_ = 156.3 mmHg) with relatively elevated diastolic stiffness (*E*_*passive*_(*V*_*peak*_) = 0.81 mmHg*ml^-1^) and a modest contractile reserve (*PVAi*_*is*o_ = 606.7 mmHg*ml*m^-2^). Patient 3 demonstrated the least dilation and greatest compliance (*V*_*o*_ = 47.6 ml; *E*_*passive*_(*V*_*peak*_) = 0.24 mmHg*ml^-1^) and the largest contractile reserve (*P*_*peak*_ = 134.4 mmHg; *PVAi*_*iso*_ = 2012.3 mmHg*ml*m^-2^). Overall, the progression from a dilated, stiff ventricle with low contractile reserve (patient 1) to a smaller, more compliant ventricle with substantially higher contractile reserve (patient 3) indicates that native LV function is most impaired in patient 1 and most preserved in patient 3. The PV envelopes reflect this pattern through a progressive leftward shift and enlargement when ordered from patient 1 to patient 3 (Fig. 3).

### 3.2 Digital Twin vs Clinical Ramp Hemodynamics

Speed-specific digital twins reproduced clinical ramp hemodynamics with percent residuals of ≤ 20% across all variables and pump speeds, except for a single CVP residual of 33.3% at 5,600 RPM in patient 1, which corresponded to an absolute error of 1 mmHg (Table 3). CI residuals ranged from 0% to 18.5%, and the twins captured paradoxical CI declines with HM3 ramp-up in patient 1 (5,300→5,600 RPM) and patient 3 (5,400→5,800 RPM). The digital twins closely matched CVP in all cases, with absolute errors ≤ 3 mmHg, even when low reference pressures amplified percent residuals. AP and PAP residuals remained < 20% across all speeds and were typically ≤ 15%, with exact matches in several instances. PAWP exhibited the largest relative deviations, with residuals up to 20% (patients 1 at 5,900 RPM and 2 at 5,800 RPM). Notably, the 5300 RPM condition in patient 1, which was deliberately withheld from the parameter identification process, was reproduced with ≤ 10% residuals across reported outputs. These results indicate that the personalized RDEF can be generalized to pump speeds not utilized for its parameterization with high fidelity, supporting its robustness and clinical usefulness.

**Table 3.** Summary of speed-specific digital twin ramp hemodynamics for the HM3 patient cohort. The % residuals between clinical and digital twin ramp hemodynamics are included. ^†^% residual not reported at 5900 RPM for patient 1’s PAWP due to a clinical reference value of 0mmHg.

| Patient ID | Speed (RPM) | CI (l*min<sup>-1</sup>*m<sup>-2</sup>) | CVP (mmHg) | PAWP (mmHg) | AP (mmHg) | PAP (mmHg) |
| --- | --- | --- | --- | --- | --- | --- |
| 1 | 5000 | 1.45<br>(3.3%) | 11<br>(0.0%) | 22<br>(8.3%) | 105/82 (89)<br>[4.0% / 5.1% (3.5%)] | 38/23 (31)<br>[5.6% / 0.0% (3.1%)] |
|  | 5300 | 1.73<br>(2.8%) | 3<br>(0.0%) | 19<br>(9.5%) | 114/88 (95)<br>[8.8% / 7.3% (2.2%)] | 35/21 (25)<br>[5.4% / 4.5% (7.4%)] |
|  | 5600 | 1.64<br>(0.0%) | 4<br>(33.3%) | 19<br>(0.0%) | 114/99 (103)<br>[0.9% / 2.1% (0.0%)] | 39/19 (26)<br>[11.4% / 11.8% (0.0%)] |
|  | 5900 | 2.20<br>(9.3%) | 1<sup>+</sup> | 12<br>(20.0%) | 106/62 (90)<br>[6.2% / 6.9% (9.8%)] | 34/16 (21)<br>[9.7% / 14.3% (8.7%)] |
| 2 | 5100 | 1.76<br>(1.6%) | 7<br>(12.5%) | 16<br>(11.1%) | 125/79 (89)<br>[5.9% / 0.0% (2.2%)] | 52/24 (33)<br>[10.3% / 14.3% (8.3%)] |
|  | 5400 | 1.91<br>(0.0%) | 8<br>(0.0%) | 12<br>(7.7%) | 143/67 (90)<br>[1.4% / 15.5% (1.1%)] | 49/24 (33)<br>[2.0% / 0.0% (2.9%)] |
|  | 5800 | 2.00<br>(4.3%) | 8<br>(0.0%) | 12<br>(20.0%) | 122/88 (99)<br>[1.6% / 0.0% (3.9%)] | 47/22 (29)<br>[6.0% / 12.0% (3.6%)] |
| 3 | 5000 | 1.70<br>(6.1%) | 15<br>(6.3%) | 17<br>(10.5%) | 117/77 (86)<br>[17.0% / 6.9% (4.9%)] | 35/23 (25)<br>[0.0% / 4.2% (7.4%)] |
|  | 5400 | 2.1<br>(18.5%) | 19<br>(18.8%) | 13<br>(13.3%) | 132/44 (90)<br>[0.0% / 17.0% (18.4%)] | 37/22 (25)<br>[5.1% / 0.0% (0.0%)] |
|  | 5800 | 1.73<br>(0.0%) | 15<br>(6.3%) | 11<br>(0.0%) | 129/96 (103)<br>[5.7% / 4.3% (0.0%)] | 37/21 (24)<br>[8.8% / 5.0% (0.0%)] |

## 4. Discussion

This study advances LVAD-supported physiology modeling by introducing the RDEF model and demonstrating its feasibility within a hybrid verification environment. The RDEF model constructs a patient-specific PV envelope from routine ramp hemodynamics by parameterizing a monotonic pPVR and a unimodal aPVR (Fig. 3; Table 2), then combines these relations via weighted superposition to simulate beat-to-beat PV dynamics (Eq. 3). Embedding the personalized RDEF within PSCOPE digital twins reproduced patient ramp responses, thereby supporting its PV envelope representation and ramp-based parameter identification strategy. These findings establish the RDEF model as a tractable, mechanistically interpretable representation of HM3-assisted LV function, enabling physiologically faithful PSCOPE digital twins without requiring invasive PV-loop acquisition.

PSCOPE-based digital twins reproduced patient-specific preload and afterload sensitivities of HM3-assisted LV function across ramp speeds (Table 3). In this context, the ventriculo-arterial coupling between the unimodal aPVR and arterial load regulates afterload sensitivity. Critically, the unimodal form permits LV pressure to decline along the descending Frank–Starling limb characteristic of dilated failure. The monotonic pPVR captures preload sensitivity by encoding diastolic stiffness and mapping pulmonary venous return to LV filling pressures across speeds. Within the digital-twin environment, the closed-loop interaction between experimental–numerical loading conditions and the personalized PV relations yielded residuals that generally remained ≤ 20% for CI, AP, PAP, PAWP, and CVP, which is comparable to reported inter-test variability and measurement error in clinically acquired hemodynamics [45-47]. This level of agreement supports the use of PSCOPE digital twins as faithful approximations of in vivo HM3–LV mechanics under clinically representative loading conditions.

Beyond its clinical motivation, RDEF contributes a ventricular modeling approach that improves upon common LV representations that impose monotonic or near-linear aPVRs and therefore cannot capture the descending Frank–Starling limb. Such simplifications can mischaracterize LV adaptations during ramp unloading [48-50]. To address this shortcoming, Wang et al. introduced a unimodal aPVR but implemented it as a piecewise bilinear approximation for tractability [29]. Although this approach preserves a single peak, it introduces a discontinuity in systolic elastance at *V*_*peak*_, promotes supraphysiologic pressure estimates, and presents practical challenges for patient-specific identification because its piecewise structure requires estimating multiple parameters per segment from ramp data that may not fully constrain them. In contrast, we employ an exponential unimodal aPVR that allows systolic elastance to vary continuously with LV unloading, thereby better reflecting the pathophysiology of failing ventricles (Fig. 1).

We generated patient-specific PV envelopes by fitting the pPVR and aPVR using speed-specific (LVEDV, PAWP) and (LVEDV, sAP) targets, respectively. The resulting envelope provides a relatively load-insensitive characterization of intrinsic LV mechanical potential by relating chamber volume to pressures the myocardium can sustain passively and generate actively. Routine ramp measurements identify the envelope’s patient-specific parameter values, enabling clinicians to infer the PV behavior of LVAD-assisted ventricles without invasive instrumentation. The PV envelope can be reported as a standalone construct or embedded within PSCOPE via the RDEF model for closed-loop verification against physical LVAD dynamics.

The pPVR parameters {*A, B, V*_*o*_} characterize passive diastolic mechanics: *A* sets the pressure scale, *B* governs how stiffness escalates with filling, and *V*_*o*_ shifts the PV envelope along the volume-axis. The reference elastance *E*_*passive*_(*V*_*peak*_) offers a compact index for longitudinal tracking of diastolic dysfunction. Clinically, increases in *E*_*passive*_(*V*_*peak*_) indicate progressive diastolic stiffening consistent with hypertrophy, whereas rightward shifts in *V*_*o*_ indicate adverse chamber dilation. The aPVR parameters {P_*peak*_,*V*_*peak*_, ΔV_*peak*_}, with ΔV_*peak*_ = V_*peak*_ − V_*o*_, quantify active contractile capacity: *P*_*peak*_ quantifies maximal pressure generation, V_*peak*_ identifies the volume at which the LV attains P_*peak*_, and ΔV_*peak*_ specifies the volume range for increasing systolic function; beyond this range, the LV operates under declining systolic function [34]. These parameters distinguish ventricles with limited function from those that tolerate broader load variation, and yield the energetic index *PVAi*_*is*o_, which quantifies contractile reserve at peak systolic function. Collectively, the physiologic content of this parameterization supports longitudinal monitoring of LV remodeling for optimal LVAD management.

Under continuous-flow unloading, state-dependent uncertainties affect the reliability of the surrogate operating points (LVEDV, PAWP) and (LVEDV, sAP). At high speeds with aortic valve closure, LV systolic pressure falls below aortic systolic pressure (so sAP overestimates LV systolic pressure), while LVEDV reasonably approximates LVESV [28-29]. At lower speeds with frequent aortic valve opening, sAP more closely reflects LV systolic pressure, but LVESV diverges from LVEDV. We therefore recommend sampling at least three ramp speeds with echocardiographically distinct aortic valve behavior—a high-speed condition with valve closure, a low-speed condition with beat-to-beat opening, and an intermediate setting with intermittent opening—to anchor both limbs of the aPVR and mitigate opposing pressure and volume biases. For the diastolic surrogate, the approximation PAWP ≈ LV end-diastolic pressure is most reliable at hemodynamic steady states without substantial mitral regurgitation, interventricular septal shift, arrhythmia, or large respiratory swings. Clinicians should average PAWP over multiple beats and acquire it at settings that promote favorable valvular and septal dynamics. Maximizing PV envelope fidelity may require sampling end-systolic and end-diastolic surrogates at different speeds within a given ramp session, reflecting state-dependent trade-offs in pressure and volume uncertainties during ramp unloading.

Finally, the retrospective single-center design and small cohort (n=3) constrain the generalizability of the findings in this study. In addition, LVEDV was derived from LVEDD using the Teichholz relation (Eq. 6), which is susceptible to geometric error in asymmetric ventricles. Future work should focus on larger, multicenter longitudinal studies and prioritize ramp protocols that explicitly mitigate surrogate uncertainty to strengthen the parameterization and validation of the PV envelope.

## 5. Conclusion

This study characterized HM3-assisted LV mechanics using the novel RDEF model, which combines a monotonic pPVR and a unimodal aPVR to define a PV envelope that represents LV behavior during diastolic filling and peak systolic contraction. Ramp-derived end-diastolic (LVEDV, PAWP) and end-systolic (LVEDV, sAP) surrogate pairs provided fitting targets for patient-specific parameterization of the PV envelope, and PSCOPE digital twins incorporated the fitted envelope via a weighted superposition to reproduce each patient’s ramp hemodynamics. Across patients and pump speeds, these digital twins reproduced clinical hemodynamics with residuals of ≤ 20% for CI, PAWP, AP, and PAP (Table 3). The digital twins matched CVP within ≤3 mmHg in all cases, although one setting produced a 33.3% residual due to a low reference pressure. For patient 1, the 5300 RPM speed deliberately withheld from PV calibration was reproduced with residuals ≤ 10% across reported outputs, indicating that the personalized RDEF can support accurate PSCOPE digital twin modeling at pump speeds beyond those used for its parameter identification.

The PV envelopes convey clinically meaningful heterogeneity through key geometric features such as the volume-axis intercept, peak active pressure, and enclosed area. Together, these features compactly summarize diastolic stiffness, the volume range for increasing systolic function, maximal contractile capability, and contractile reserve. These mechanistic descriptors complement conventional load-dependent indices and can illuminate inter-patient differences that standard clinical measurements may not distinguish.

Clinicians can report the PV envelope as a standalone construct or embed its parameters within PSCOPE for closed-loop verification against physical LVAD fluid dynamics, thereby balancing deployability with physiological rigor. Their clinical utility, however, is contingent on the state-dependent fidelity of the surrogate operating points (LVEDV, PAWP) and (LVEDV, sAP). Ramp protocols should therefore adopt sampling strategies that mitigate opposing pressure and volume uncertainties and explicitly account for valvular and septal dynamics during unloading.

Overall, these results support the RDEF model as a mechanistically interpretable and clinically tractable representation of native LV function under HM3 support. Prospective, multicenter studies are now warranted to confirm whether envelope parameters enable reliable longitudinal assessment of myocardial recovery.

## Notes

### Competing Interest Statement

Author Arman Kilic is a consultant and speaker for Abbott, Abiomed, 3ive, and LivaNova. The remaining authors declare that the study was conducted without any financial or personal interests that could be construed as potential conflicts of interest.

